# The joint evolution of animal movement and competition strategies

**DOI:** 10.1101/2021.07.19.452886

**Authors:** Pratik R. Gupte, Christoph F. G. Netz, Franz J. Weissing

**Affiliations:** Groningen Institute for Evolutionary Life Sciences, University of Groningen, Groningen 9747 AG, The Netherlands

**Keywords:** Movement ecology, Intraspecific competition, Individual differences, Foraging ecology, Kleptoparasitism, Individual based modelling, Ideal Free Distribution

## Abstract

Competition typically takes place in a spatial context, but eco-evolutionary models rarely address the joint evolution of movement and competition strategies. Here we investigate a spatially explicit producer-scrounger model where consumers can either forage on a heterogeneous resource landscape or steal resource items from conspecifics (kleptoparasitism). We consider three scenarios: (1) a population of foragers in the absence of kleptoparasites; (2) a population of consumers that are either specialized on foraging or on kleptoparasitism; and (3) a population of individuals that can fine-tune their behavior by switching between foraging and kleptoparasitism depending on local conditions. By means of individual-based simulations, we study the joint evolution of movement and competition strategies, and we investigate the implications on the resource landscape and the distribution of consumers over this landscape. In all scenarios and for all parameters considered, movement and competition strategies evolved rapidly and consistently across replicate simulations. The evolved movement and resource exploitation patterns differ considerably across the three scenarios. For example, foragers are attracted by conspecifics in scenario (1), while they are repelled by conspecifics in scenario (2). Generally the movement strategies of kleptoparasites differ markedly from those of foragers, but even within each class of consumers polymorphisms emerge, corresponding to pronounced differences in movement patterns. In all scenarios, the distribution of consumers over resources differs substantially from ‘ideal free’ predictions. We show that this is related to the intrinsic difficulty of moving effectively on a depleted landscape with few reliable cues for movement. Our study emphasises the advantages of a mechanistic approach when studying competition in a spatial context, and suggests how evolutionary modelling can be integrated with current work in animal movement ecology.

## Introduction

Intraspecific competition is an important driver of population dynamics and the spatial distribution of organisms (Krebs and Davies, 1978), and can be broadly classified into two main types, ‘exploitation’ and ‘interference’. In exploitation competition, individuals compete indirectly by depleting a common resource, while in interference competition, individuals compete directly by interacting with each other (Birch, 1957; Case and Gilpin, 1974; Keddy, 2001). A special case of interference competition which is widespread among animal taxa is ‘kleptoparasitism’, in which an individual steals a resource from its owner (Iyengar, 2008). Since competition has an obvious spatial context, animals should account for the locations of intraspecific foraging competitors when deciding where to move (Nathan et al., 2008). Experimental work shows that indeed, competition, as well as the pre-emptive avoidance of competitive interactions, affects animal movement decisions in taxa as far apart as waders (Goss-Custard, 1980; Vahl et al., 2005*a*; Rutten et al., 2010*b*, see also Rutten et al. 2010*a*; Bijleveld et al. 2012), and fish (Laskowski and Bell, 2013). This is expected to have downstream effects on animal distributions at relatively small scales, such as across resource patches (see Fretwell and Lucas, 1970), as well as at larger scales, determining species distributions (e.g. Duckworth and Badyaev, 2007, see Schlägel et al. 2020 for background). Animal movement decisions are thus likely to be adaptive responses to landscapes of competition, with competitive strategies themselves being evolved responses to animal distributions. Studying this joint evolution is key to understanding the spatial distribution of animals, but empirical studies are nearly impossible at large spatio-temporal scales. This makes models linking individual traits and behavioural decisions to population distributions necessary.

Contemporary individual-to-population models of animal space-use (reviewed in DeAngelis and Diaz, 2019) and competition, however, are only sufficient to represent very simple movement and prey-choice decisions, and struggle to adequately represent more complex systems of consumer-resource interactions. For example, models including the ideal free distribution (IFD; Fretwell and Lucas, 1970), information-sharing models (Giraldeau and Beauchamp, 1999; Folmer et al., 2012), and producer-scrounger models (Barnard and Sibly, 1981; Vickery et al., 1991; Beauchamp, 2008), often treat foraging competition in highly simplified ways. Most IFD models, for instance, consider resource depletion unimportant or negligible (continuous input models, see Tregenza, 1995; van der Meer and Ens, 1997), or make simplifying assumptions about interference competition, even modelling an *ad hoc* benefit of grouping (e.g. Amano et al., 2006). Producer-scrounger models primarily examine the benefits of choosing either a producer or scrounger strategy given local conditions, such as the number of conspecifics (Vickery et al., 1991), or the order of arrival on a patch (Beauchamp, 2008). Moreover, these models simplify the mechanisms by which competitive decisions are made, often ignoring spatial structure (see also Holmgren, 1995; Garay et al., 2020; Spencer and Broom, 2018).

On the contrary, competition occurs in a spatial context, and spatial structure is key to foraging (competition) decisions (Beauchamp, 2008). Consequently, the abundance of resources and their depletion, as well as the presence of potential competitors is of obvious importance to individuals’ movement decisions (resource selection, *sensu* Manly et al., 2007). How animals are assumed to integrate the costs (and potential benefits) of competition into their movement decisions has important consequences for theoretical expectations of population distributions (van der Meer and Ens, 1997; Hamilton, 2002; Beauchamp, 2008). In addition to short-term, ecological effects, competition should also have evolutionary consequences for individual *movement strategies*, as it does for so many other aspects of behaviour (Baldauf et al., 2014), setting up feedback loops between ecology and evolution. Modelling competition and movement decisions jointly is thus a major challenge. A number of models take an entirely ecological view, assuming that individuals move or compete ideally, or according to some fixed strategies (Vickery et al., 1991; Holmgren, 1995; Tregenza, 1995; Amano et al., 2006, but see Hamilton 2002). Models that include evolutionary dynamics in the movement (de Jager et al., 2011, 2020) and foraging competition strategies (Beauchamp, 2008; Tania et al., 2012) are more plausible, but they too make arbitrary assumptions about the functional importance of environmental cues to individual decisions.

Furthermore, populations likely contain significant individual variation in movement and competition characteristics, such that individuals make different decisions given similar cues (Laskowski and Bell, 2013). Capturing these differences in models is likely key to better understanding how individual decisions scale to population- and community-level outcomes (Bolnick et al., 2011). Individual based models are well suited to capturing variation in responses to environmental cues, and also force researchers to be explicit about their modelling assumptions, such as *how exactly* competition affects fitness. Similarly, rather than taking a purely ecological approach and assuming such differences (e.g. in movement rules White et al., 2018), modelling the evolution of movement strategies in a competitive landscape can reveal whether individual variation emerges in plausible ecological scenarios (as in Getz et al., 2015). This allows the functional importance of environmental cues to movement and competition decisions in evolutionary models to be joint outcomes of selection, leading, for example, different competition strategies to be associated with different movement rules (Getz et al., 2015).

Here, we present a mechanistic, model of intraspecific foraging competition in a spatially explicit context, where competition is shaped by the joint evolution of foraging competition and movement strategies. As foraging and movement decisions are taken by individuals, we study the joint evolution of both types of decision-making by means of an individual-based simulation model. Such models are well suited to modelling the ecology and evolution of complex behaviours (Guttal and Couzin, 2010; Kuijper et al., 2012; Getz et al., 2015, 2016; White et al., 2018; Long and Weissing, 2020; Netz et al., 2020, for conceptual underpinnings see Huston et al. (1988); DeAngelis and Diaz (2019)). This allows us to both focus more closely on the interplay of exploitation and interference competition, and to examine the feedback between movement and foraging behaviour at ecological and evolutionary timescales. In our model, foraging individuals move on a spatially fine-grained resource landscape with discrete, depletable food items that need to be processed (‘handled’) before consumption. Foragers make movement decisions using an inherited (and evolvable) strategy which integrates local cues, such as the local resource and competitor densities. After each move, individuals choose between two foraging strategies: whether to search for a food item or steal from another individual; the mechanism underlying this foraging choice is also inherited. We take lifetime resource consumption as a proxy for fitness, such that more successful individuals produce more offspring, and thus are more successful in transmitting their movement and foraging strategies to future generations (subject to small mutations). We consider three scenarios: in the first scenario, we examine only exploitation competition. In the second scenario, we introduce kleptoparasitic interference as an inherited strategy, fixed through an individual’s life. In the third scenario, we model kleptoparasitism as a behavioural strategy conditioned on local environmental and social cues.

Our model allows us to examine the evolution of individual movement strategies, population-level resource intake, and the spatial structure of the resource landscape. The model enables us to take ecological snapshots of consumer-resource dynamics (animal movement, resource depletion, and competition) proceeding at evolutionary time-scales. Studying these snapshots from all three scenarios allows us to check whether, when, and to what extent the spatial distribution of competitors resulting from the co-evolution of competition and movement strategies corresponds to standard IFD predictions. We investigate three primary questions: *(1)* What movement patterns will evolve in producer-scrounger systems? To what extent will the pattern differ between producers and scroungers? *(2)* Does the (evolved) spatial distribution of consumers and resources correspond to “ideal free” expectations? To what extent is the outcome dependent on the modeling scenarios considered? *(3)* Do individuals in the same “competition” state use the same movement strategy or are there indications for systematic individual differences in movement patterns?

## The Model

Individual-based models have the advantage and the disadvantage that they have to explicitly specify numerous assumptions (e.g. on the spatial structure, the interaction structure, the timing of events), while the same kind of assumptions are often hidden below the surface in analytical models. As we are mainly interested in general, conceptual insights, we tried to keep our model assumptions as simple and generic as possible. However, to keep the model realistic (and to relate model outcomes with empirical observations) the model set-up is inspired by the foraging behavior of shorebirds *Charadrii*. This is reflected by the gridded structure of the environment, the capacity of each grid cell to hold multiple individuals, the discrete nature of the resources, and the discrete conception of time within and between generations. Shorebirds such as oyster-catchers (*Haematopus* spp.) are a convenient model system, and are extensively studied in the context of foraging competition, both empirically (e.g. Vahl et al., 2005*a*,*b*, 2007; Rutten et al., 2010*a*,*b*), and using individual-based models (reviewed in Stillman and Goss-Custard, 2010). We simulated a population with a fixed number of individuals (N = 10,000), which move on a landscape of 512^2^ grid cells (approx. 1 individual per 26 cells), with wrapped boundaries; individuals passing beyond the bounds at one end re-appear on the opposite side. The model has two time scales, first, an ecological time scale of *T* timesteps comprising one generation (default T = 400), during which individuals move, make foraging decisions, and handle prey-items they find or steal. Individuals are immobile while handling food items, creating the conditions for kleptoparasitism (Brockmann and Barnard, 1979; Ruxton et al., 1992). On the second, evolutionary time scale of 1,000 generations, individuals reproduce, transmitting their movement and foraging strategies to their offspring, whose number is proportional to individual intake at the ecological time scale.

### Resource Landscape

#### Prey Abundance

We considered a resource landscape that is heterogeneous in its productivity of discrete resources, but with strong spatial clustering of grid cells of similar productivity. We considered our discrete resources, called ‘prey-items’ to represent mussels, a common prey of many shorebirds, whose abundances are largely driven by external gradients. We assigned each cell a constant probability of generating a new prey item per timestep, which we refer to as the cell-specific growth rate *r*. We modelled clustering in landscape productivity by having the distribution of *r* across the grid take the form of 1,024 resource peaks, placed at regular distances of 16 grid cells from the peaks around them; *r* declines from the centre of each peak (called *r*_*max*_) to its periphery (see Fig. 1C). Thus the central cell generates prey-items five times more frequently than peripheral cell: at *r*_*max*_ = 0.01, central cells generate one item per 100 timesteps (four items/generation), while the peripheral cells generate one item only every 500 timesteps (*<* one item/generation). All landscape cells have a uniform carrying capacity *K* of 5 prey-items.

**Figure 1:**
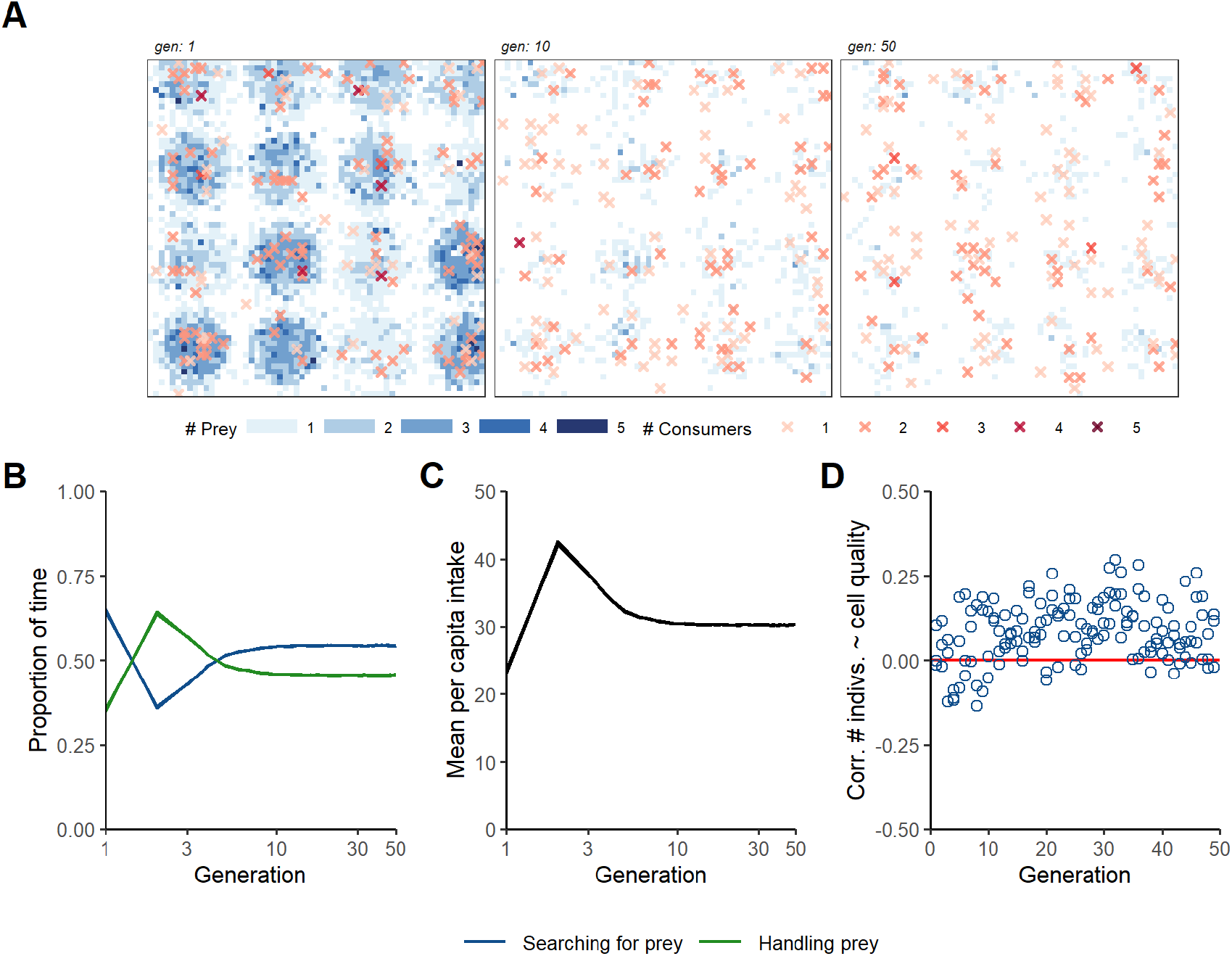
Eco-evolutionary implications of pure exploitation competition (scenario 1). **(A)** When a population is comprised solely of foragers seeking prey on a resource landscape, the initially well-stocked resource landscape is swiftly depleted within 10 generations (out of 1,000 simulated). This sparsity in prey-item abundance is maintained throughout the remaining generations of the simulation. Individuals, whose local density is shown by coloured crosses, are scattered over the landscape. These dynamics are explained by the fact that **(B)** within 20 generations of evolution, the population reaches an equilibrium in the relative proportion of time spent on searching prey and handling prey, and in **(C)** the total intake of the population. **(D)** In a departure from the intake matching rule of IFD theory, cell occupancy (number of foragers per cell) is only weakly correlated with cell productivity *r*. Panel **A** shows a single replicate, while panels **B, C** and **D** show three replicate simulations (lines overlap almost perfectly); all panels are for *r*_*max*_ = 0.01. NB: Both **B, C** show a log-scaled X axis to more clearly show dynamics in early generations.

#### Prey Acquisition by Foragers

Foragers perceive a cue indicating the number of prey-items *P* in a cell, but fail to detect each item with a probability *q*, and are thus successful in finding a prey-item with a probability 1 − *q*^*P*^. Individuals on a cell forage in a randomised sequence, and the probability of finding a prey-item (1 − *q*^*P*^) is updated as individuals find prey, reducing *P*. Foragers that find a prey-item must handle it for a fixed handling time *T*_*H*_ (default = 5 timesteps), before consuming it (Ruxton et al., 1992). Natural examples include the time required for an oystercatcher to break through a mussel shell, or a raptor to subdue prey; overall, the handling action is obvious, and the prey is not fully under the control of the finder (Brockmann and Barnard, 1979). Foragers that do not find a prey-item are considered idle in that timestep, and are counted as ‘non-handlers’. Similarly, handlers that finish processing their prey in timestep *t* can only forage again in timestep *t* + 1, i.e., they are idle in the timestep *t*.

### Movement and Competition Strategies

#### Movement Strategies

We model movement as comprised of small, discrete steps of fixed size, which are the outcome of individual movement decisions made using evolved movement strategies. Across scenarios, individuals make movement decisions by selecting a destination cell, after assessing potential destinations based on available cues (similar to step selection or resource selection; Fortin et al., 2005; Manly et al., 2007), and similar to the approach used previously by Getz et al. (2015, 2016) and White et al. (2018). At the end of each timestep *t*, individuals scan the nine cells of their Moore neighbourhood for three environmental cues, *(1)* an indication of the number of discrete prey items *P, (2)* the number of individuals handling prey *H* (referred to as ‘handlers’), and *(3)* the number of individuals not handling prey *N* (referred to as ‘non-handlers’). Individuals rank potential destinations (including the current cell) by their suitability *S*, where *S* = *s*_*P*_*P* + *s*_*H*_*H* + *s*_*N*_*N*, and move to the most suitable cell in timestep *t* + 1. The weighing factors for each cue, *s*_*P*_, *s*_*H*_, and *s*_*N*_, are evolvable traits, and are genetically encoded and transmitted between generations. All individuals move simultaneously, and then implement their foraging or kleptoparasitic behaviour to acquire prey. However, handlers do not make any movements until they have fully handled and consumed their prey.

#### Scenario 1: Exploitative Competition

In scenario 1, we simulate only exploitative competition; individuals (henceforth called ‘foragers’) move about on the landscape and probabilistically find, handle, and consume prey items. Foragers can be either in a ‘searching’ or a ‘handling’ state (Holmgren, 1995). The only evolvable properties are the cue weighing factors which determine the suitability scores (*s*_*P*_, *s*_*H*_ and *s*_*N*_).

#### Scenario 2: Fixed Interference Competition

In scenario 2, the competition strategy is genetically determined and transmitted from parents to offspring: exploitative competition (by foragers), or kleptoparasitic interference (by kleptoparasites). Each of these strategies can evolve a (separate) movement strategy. Kleptoparasites cannot extract prey-items directly from the landscape, and only steal from handlers (see Holmgren, 1995). Kleptoparasites are modelled as always being successful in stealing from handlers, and such successful surprise attacks are commonly observed among birds (Brockmann and Barnard, 1979). However, if multiple kleptoparasites target the same handler, only one (randomly selected) is considered successful — thus kleptoparasites compete exploitatively among themselves. Handlers robbed of prey subsequently ‘flee’ up to 5 cells away from their location. Having acquired prey, kleptoparasites become handlers, but need only handle prey for *T*_*H*_ − *t*_*h*_ timesteps, where *t*_*h*_ is the time that the prey has already been handled by its previous owner. Unsuccessful kleptoparasites are considered idle, and are counted as non-handlers.

#### Scenario 3: Conditional Interference Competition

In scenario 3, each individual can either act as a forager, or as a kleptoparasite, depending on its assessment of local circumstances. Individuals process the cell-specific environmental cues *P, H*, and *N* to determine their location in the next timestep (based on their inherited movement strategy). Additionally, individuals process cell-specific environmental cues in timestep *t* to determine their strategy in the next timestep as

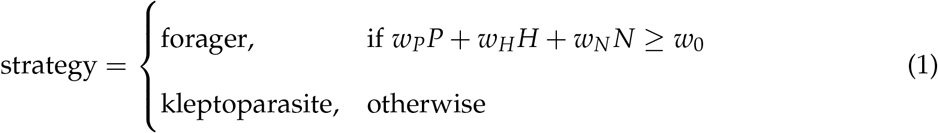

where the cue weights *w*_*P*_, *w*_*H*_ and *w*_*N*_, and the threshold value *w*_0_, are also heritable between generations. Apart from the ability to switch between foraging and kleptoparasitism, the competition dynamics are the same as in scenario 2.

### Reproduction and Inheritance

Our model considers a population of fixed size (10,000 individuals) with discrete, non-overlapping generations. Individuals are haploid and reproduction is asexual. Each individual has 7 gene loci that encode the decision making weights; only the weights in control of individual movement (*s*_*P*_, *s*_*H*_, *s*_*N*_) are active in scenarios 1 and 2. In scenario 3, the weights for foraging decisions (*w*_*P*_, *w*_*H*_, *w*_*N*_, *w*_0_) are also active, and are transmitted from parent individuals to offspring. Hence the alleles at these loci correspond to real numbers that are transmitted from parent individuals to their offspring.

Each individual’s number of offspring is proportional to the individual’s total lifetime intake of resources; hence, resource intake is used as a proxy for fitness. A weighted lottery (with weights proportional to lifetime resource intake) selects a parent for each offspring in the subsequent generation (prior implementation in Tania et al., 2012; Netz et al., 2020). Across scenarios, the movement decision-making weights are subject to rare, independent mutations (*µ* = 0.001). The mutational step size (either positive or negative) is drawn from a Cauchy distribution with a scale of 0.01 centred on zero, allowing for a small number of very large mutations while most mutations are small. In scenarios 1 and 2, the foraging-decision weights are not relevant. However, in scenario 2, we allow a forager to infrequently mutate into a kleptoparasite (or *vice versa*; *µ* = 0.001). In scenario 3, the foraging weights also mutate as described above. We initialised each offspring at random locations on the landscape, leading individuals to experience conditions potentially very different from those of their parent.

### Simulation Output and Analysis

We ran all three scenarios at a default *r*_*max*_ of 0.01, which we present in the Results, and also across a range of *r*_*max*_ values between 0.001 and 0.05 (see Fig. 6 and Supplementary Material Figs. – 1.3). We initialised the decision making weights with values uniformly distributed between -1.0 and 1.0, to allow sufficient variation in the population.

#### Population Activities and Intake

Across scenarios, in each generation, we counted the number of times foragers were searching for prey, kleptoparasites were searching for handlers, and the number of timesteps that individuals of either strategy were handling a prey-item. We refer to the ratio of these values as the population’s ‘activity budget’. We examined how the population activity budget developed over evolutionary time, and whether a stable equilibrium was reached. Furthermore, we counted the population’s mean per-capita intake per generation as a measure of population productivity.

#### Visualising Decision-Making Weights

To understand the evolution of individual movement and competition strategies, we exported the decision-making weights of each individual in every generation of the simulation. To visualise functional differences in weights, which could take arbitrarily large values, we multiplied each weight by 20 and applied a hyperbolic tangent transform. This scaled the weights between -1 and +1, and we plotted these weights to understand individual variation in movement rules, as well as calculating how preference and avoidance of cues evolved across scenarios.

#### Ecological Snapshots of Consumer-Resource Distributions

We exported snapshots of the entire simulation landscape at the mid-point of each generation (*t* = 200). Each snapshot contained data on *(1)* the number of prey-items, *(2)* the number of handling individuals, and the number of individuals using either a *(3)* searching forager strategy or *(4)* kleptoparasitic strategy, on each cell. We used a subset of the total landscape (60^2^ of 512^2^ cells) for further analyses to speed up computation. We determined the availability of direct resource cues for movement in each cell by calculating the cell-specific item gradient for each landscape snapshot, as the difference in prey counts between each cell and its neighbouring cells. For each generation, we calculated the proportion of cells from which it was possible to sense differences in prey-items, i.e., a neighbouring cell with either more or fewer items.

#### Testing the Input Matching Rule

A basic prediction of the IFD and the related matching rule is that the number of individuals on occupied patches should be proportional to patch productivity (Fretwell and Lucas, 1970; Parker, 1978; Houston, 2008). Patch productivity is challenging to measure in real world systems, but is among our model’s building blocks, and we examined the correlation between the number of individuals (excluding handlers) and the cell-specific productivity *r*, expecting large positive values.

## Results

### Scenario 1: No Kleptoparasitism

In scenario 1, foragers deplete prey-items faster than they are replenished, drastically reducing the overall number of prey within 50 generations (Fig. 1A). The population activity budget is split between searching and handling (Fig. 1B); while handling and the mean per-capita intake are both initially low, they peak within ten generations (Fig. 1C), as individuals easily acquire prey-items from the fully stocked landscape in the first few generations. With dwindling prey-items, fewer searching foragers find prey, and handling as a share of the activity budget declines to a stable ∼ 45% within 50 generations, and mean per-capita intake also stabilises (Fig. 1C). Across generations, the correlation between the number of foragers and cell productivity is only slightly positive (Fig. 1D). This is in contrast with the perfect correspondence between resource input rate and forager density (the ‘input matching rule’), which is a defining property of the IFD (Parker, 1978; Houston, 2008). Contrary to standard IFD assumptions, foragers cannot directly “sense” the local cell productivity *r*; instead they can only use the (small) number of prey items available in a cell as a cue for local productivity (“cell quality”).

### Scenario 2: Co-existence of Foragers and Kleptoparasites

In scenario 2, with fixed foraging and kleptoparasitism allowed, the spatial distribution of prey-items at equilibrium is very different from scenario 1. Consumers graze down resource peaks until few prey-items remain on the landscape; however, within 50 generations the resource landscape recovers with prey abundances higher than in the earliest generations (Fig. 2A). This is because of the emergence of kleptoparasites (Fig. 2B): in early generations, kleptoparasites are rare, and the activity budget, the mean per-capita intake, and the distribution of consumers over the landscape, are similar to scenario 1. As resources are depleted and kleptoparasite-handler ecounters become more common than forager-prey encounters, kleptoparasitism becomes the majority strategy (a stable ∼70% of the population; see Fig. 2B), and searching for handlers to rob becomes the commonest activity. However, the high frequency of this activity and the low frequency of handling, indicate that few kleptoparasites are successful at robbing handlers.

**Figure 2:**
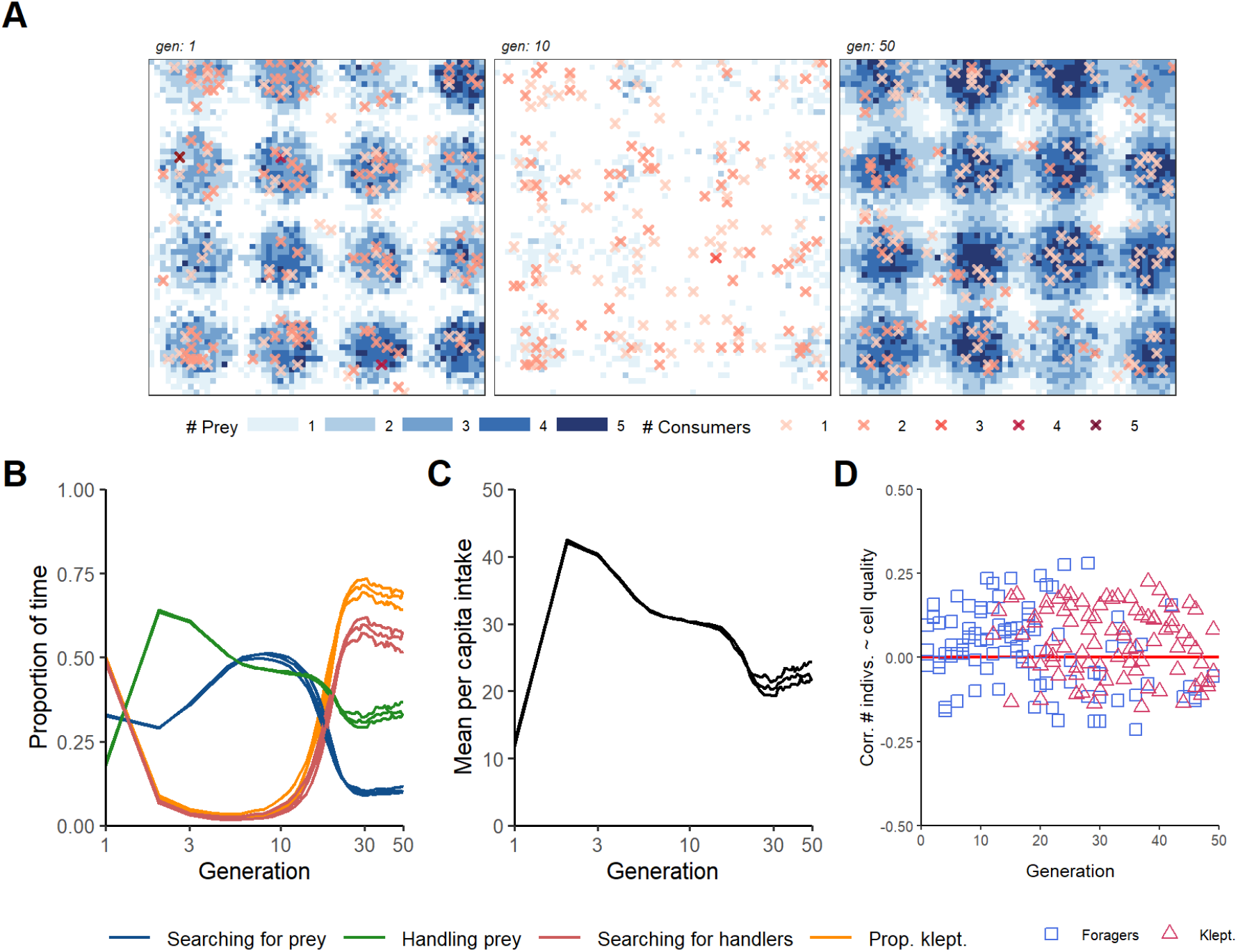
Eco-evolutionary implications of the coexistence of foragers and kleptoparasites (scenario 2). In populations with both foragers and kleptoparasites, **(A)** the initially well-stocked resource landscape is drastically depleted by generation 10; however, prey densities recover strongly by generation 50, even beyond the densities in generation 1. The local density of individuals on occupied cells is shown as coloured crosses. **(B)** An equilibrium between the strategies is reached within 30 generations, with the relative frequency of kleptoparasites (orange line) first dropping to very low levels but later recovering to reach a high level (∼70%) in all three replicates. The activity budget parallels the relative frequency of kleptoparasites, and at equilibrium, about 10% of the individuals are foragers searching for prey, 50% are kleptoparasites searching for handlers, and 40% are handlers (either foragers or kleptoparasites). **(C)** In early generations, when kleptoparasites are rare, the population intake rate exhibits the same pattern as in Fig. 1B, dropping to a lower level with the emergence of kleptoparasites. This is accompanied by an increase in the proportion of time spent on stealing attempts (red line – **B**), and a corresponding decrease in prey seeking (by searching foragers; blue line – **B**), and handling (green line – **C**). **(D)** Cell occupancy (local density of foragers per cell) is only weakly correlated with cell productivity *r*, dropping to zero at equilibrium. Panel **A** shows a single replicate, while **B, C** and **D** show three replicates; all panels are for *r*_*max*_ = 0.01.

With few foragers, few prey-items are extracted from the landscape, which recovers beyond its initial prey abundance within 50 generations (Fig. 2A). As fewer prey-items are extracted overall, mean per-capita intake also declines from an initial peak (Fig. 2C). Despite the strong spatial structure of the resource landscape within 50 generations, the correlation between consumers (of either strategy) and cell productivity remains weak or zero across generations (Fig. 2D). This may be explained by the dynamics of kleptoparasitism: foragers fleeing a kleptoparasitic attack are displaced far from their original location, and kleptoparasites must track these foragers if they are to acquire resources.

The increase of kleptoparasites from a negligible fraction to the majority strategy (Fig. 3A) is associated with an evolutionary divergence of movement strategies between foragers and kleptoparasites. While all individuals (both foragers and kleptoparasites) evolve to prefer high prey density and avoid high non-handler density (see Supplementary Material Fig. 2.2), the two types of competition strategy differ substantially in their response to handlers (Fig. 3B, 3C). Kleptoparasites very rapidly (within 3 generations) evolve a strong preference for moving towards handlers, which are their primary resource (Fig. 3B). In the absence of kleptoparasites, foragers would evolve a preference for moving towards handlers (see Supplementary Material Fig. 2.1), but, with kleptoparasites common in the population, searching foragers avoid and prefer handlers in about equal proportions (Fig. 3C). While all kleptoparasites evolve to prefer moving towards handlers, the strength of the attraction to handlers shows multiple distinct values (‘morphs’), which are remarkably persistent across generations (Fig. 3B). In replicate 3, for example, the commonest movement strategy is only weakly attracted to handlers, but this strategy coexists with various strategies that are all strongly attracted to handlers (Fig. 3B). The movement strategies of foragers show an even higher degree of polymorphism (Fig. 3C). Typically, there are no predominant movement strategies. Instead, a wide range of coexisting handler attraction/repulsion values emerges: some foragers are strongly attracted by handlers, others are strongly repelled by handlers, and yet others are neutral to the presence of handlers.

**Figure 3:**
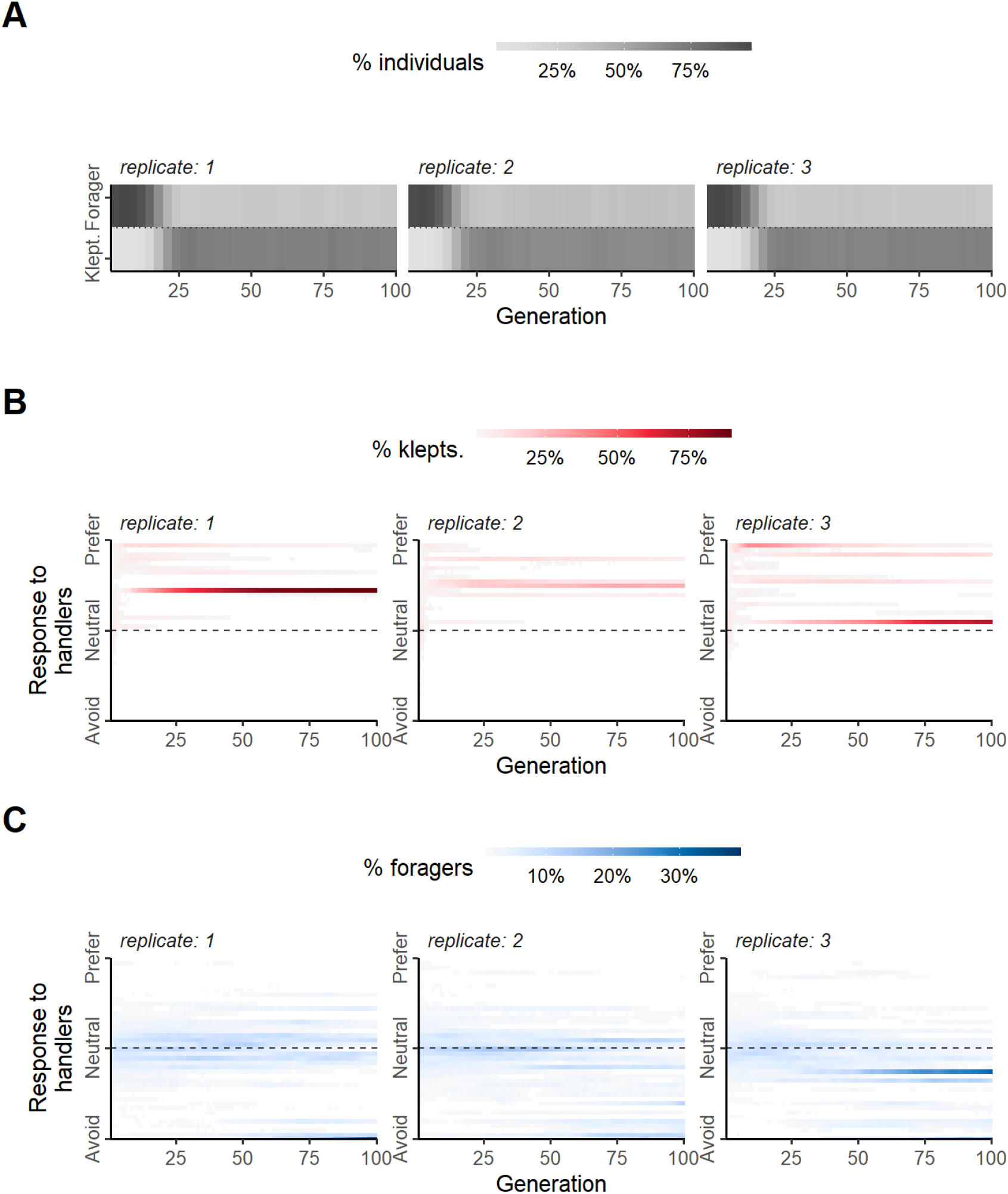
Divergence of movement strategies between foragers and kleptoparasites (scenario 2). **(A)** Kleptoparasitism rapidly becomes the more frequent strategy in scenario 2 populations for the parameters considered, with no differences across replicates. However, replicates differ considerably in the evolved movement strategies. This is illustrated by the distribution of the weighing factor *s*_*H*_ (describing the effect of local handler density on the movement decision) in kleptoparasites **(B)** and foragers **(C)**, respectively. In kleptoparasites, the weights *s*_*H*_ are generally positive, indicating that kleptoparasites are attracted by handlers. However, different *s*_*H*_ values stably coexist, indicating that kleptoparasites are polymorphic in their movement strategy. Foragers are also polymorphic in their handler responses: foragers attracted by handlers (positive *s*_*H*_) coexist with foragers repelled by handlers (negative *s*_*H*_). All panels show three replicates at *r*_*max*_ = 0.01.

### Scenario 3: Condition-dependent Kleptoparasitism

When individuals are allowed to choose their competition strategy (foraging or kleptoparasitism) based on local environmental cues, the distribution of individuals and prey items is substantially different from the two previous scenarios (Fig. 4A). Initially, as in scenario 1, individuals deplete the resource landscape of prey-items within ten generations. By generation 50, the resource landscape recovers some of the spatial structure of early generations, but prey-item abundances do not match the recovery seen in scenario 2. This too is explained by the observation that by generation 30, all individuals have a propensity to steal from handlers, i.e., when handlers are present in the vicinity, consumers will choose to target handlers for prey items, rather than forage for prey themselves (“opportunistic kleptoparasitism” ; Fig. 4B; *orange line*). However, unlike scenario 2, individuals search for prey more often and steal less (at or below 25%; compare Fig. 2B), preventing a full recovery of the resource landscape. Consequently, mean per-capita intake stabilises (after an initial spike, as in scenarios 1 and 2) within ten generations to a level similar to scenario 1 (Fig. 4C). Using conditional foraging strategies, individuals are able to switch between resource types (prey and handlers) depending on which is more profitable (Emlen, 1966), and appear to track resources. Thus, while not as strong as predicted by IFD theory, the correlations between consumer abundance and cell productivity are weakly positive (Fig. 4D).

**Figure 4:**
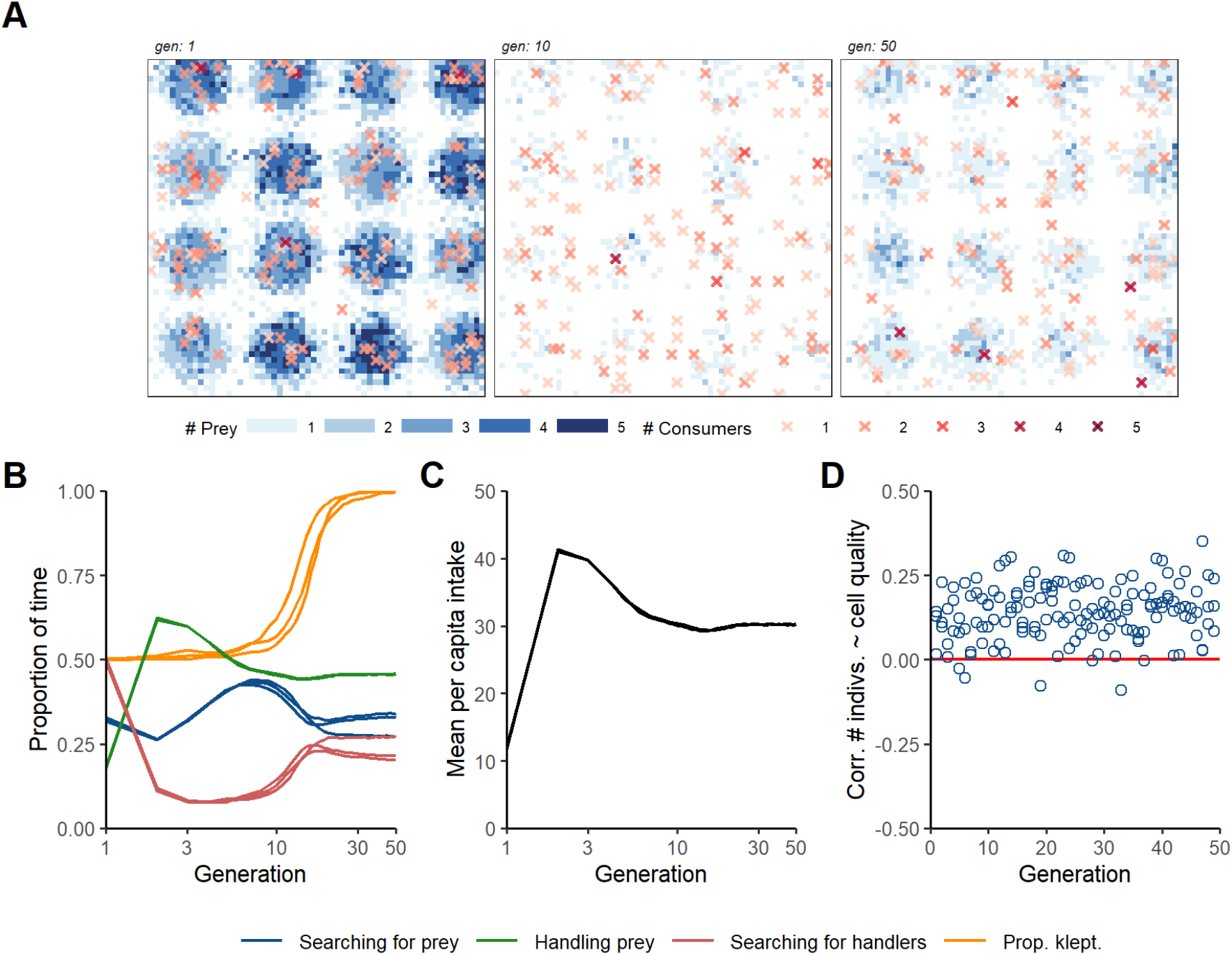
Eco-evolutionary implications of conditional foraging strategies (scenario 3). **(A)** The initially well-stocked resource landscape is rapidly depleted within 10 generations, yet within 50 generations, prey abundances recover on many cells, though not to the extent of scenario 2. The local density of individuals on occupied cells is shown as coloured crosses. **(B)** By generation 30, all individuals encountering handlers will choose to steal prey rather than search for prey themselves. The proportion of time spent searching (blue line), handling (green line), and stealing prey (red line) also reach an equilibrium that differs somewhat across replicates. **(C)** Yet, the total intake of the population reaches the same equilibrium value in all three replicates. **(D)** The correlation between the local density of individuals on a cell, and its productivity *r* is stronger than in scenario 2. Panel **A** shows a single replicate, while **B, C** and **D** show three replicates; all panels are for *r*_*max*_ = 0.01.

### Movement Rules on Depleted Landscapes

Orienting movement towards resources (Nathan et al., 2008, ; *where to move*) can be a challenge in a system with low densities of discrete prey items, because the local prey *density* may provide very limited information about local *productivity*. In our model, prey-depletion leads parts of the resource landscape to become ‘clueless regions’ (Perkins, 1992), where foragers cannot make directed movements based on prey-item abundances alone, as all neighbouring item abundances are identical (see white areas in Fig. 5A; A1: scenario 1, A2: scenario 2, A3: scenario 3). At the beginning of all three scenarios, about 75% of landscape cells have a different number of prey-items from the cells around them; these are primarily cells with an intermediate *r*, which have more prey than peripheral cells of resource peaks, but fewer prey than the central cells. This proportion rapidly declines to a much lower value within 10 generations in all three scenarios.

**Figure 5:**
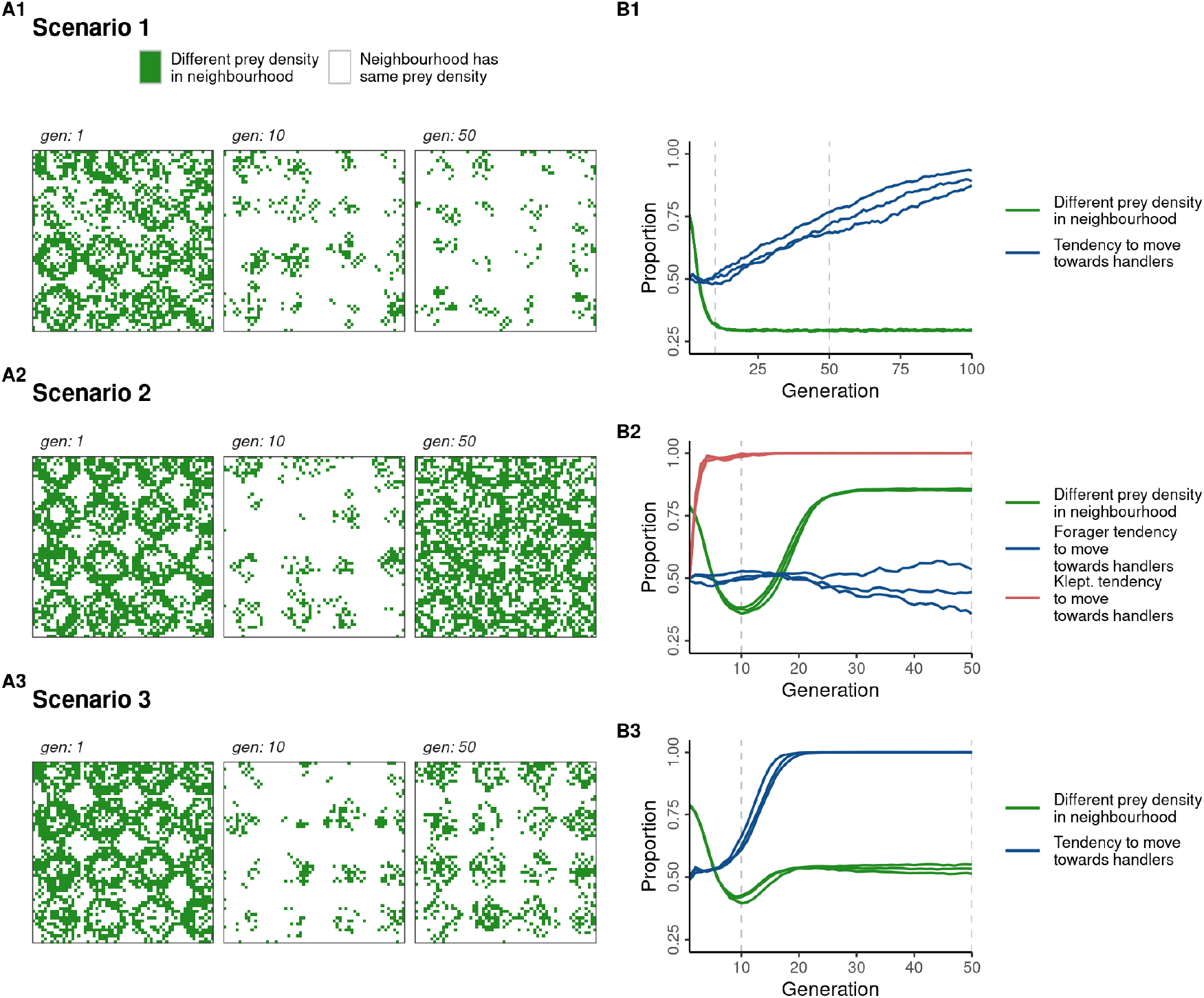
Uninformative prey densities and the evolution of alternative movement cues. **(A1, A2, A3)** On cells coloured green, local prey densities are informative for movement, as the central and neighbouring cells have different prey densities. While differences in local prey densities provide informative cues for ‘adaptive’ movement in early generations, this is much less true once the resource landscape is depleted of prey-items (depending on the scenario). **(B1, B2, B3)** The proportion of cells where differences in local prey densities provide informative movement cues (green line), and the proportion of individuals preferring to move towards handlers (blue line), whose presence may be used as an alternative cue for movement towards higher-productivity areas of the landscape. In **(B2)** representing scenario 2, this proportion is shown separately for foragers (blue line) and kleptoparasites (red line). While panels in **(A)** show a single representative replicate for *r*_*max*_ = 0.01, panels in **(B)** show three replicates.

The ‘cluelessness’ of the landscapes develops differently across scenarios on evolutionary timescales (Fig. 5B). In scenario 1, the proportion of cells with a different number of items in the neighbourhood is initially very high (Fig. 5A1). This proportion rapidly declines to ∼25% within 10 generations, as foragers deplete most prey-items, making most of the landscape a clueless region. In this context, foragers evolve to move towards handlers, with *>* 75% of individuals showing a preference for handlers within 100 generations (Fig. 5B1). Forager preference for handlers may be explained as the sensing of a long-term cue of local productivity. Since handlers are immobilised on the cell where they find a prey-item, handler density is an indirect indicator of cell *r*, and due to spatial autocorrelation, also of the *r* of bordering cells.

Scenario 2 landscapes develop similarly to scenario 1 in early generations (Fig. 5A2). However, within 50 generations, most cells bear items as extraction is reduced, with differences among cells according to their *r* (see also Fig. 2A). Thus *>* 75% of cells have a different number of items from neighbouring cells (Fig. 5A2 – panel *gen: 50*, 5B2). Unlike scenario 1, the rapid increase in handler preference is driven by kleptoparasites becoming the majority strategy (see above). Scenario 3 is similar to scenario 2, except that only about half of all cells have a different number of prey-items from neighbouring cells (Fig. 5A3, 5B3). Here, the rapid evolution of a handler preference in movement decisions cannot be assigned a clear cause, since handlers are both a potential direct resource as well as indirect cues to the location of productive cells.

### Effect of Landscape Productivity

The prey-item regrowth rate that characterises the peaks of the resource landscape (*r*_*max*_) is a measure of the productivity of the resource landscape overall. Having thus far focused on scenarios with *r*_*max*_ = 0.01 (corresponding to a peak production of 4 food times per consumer lifetime), we find that, not unexpectedly, the value of *r*_*max*_ has a marked effect on evolved population activity budgets, mean per capita intake, and even evolved strategies. The frequency of foraging reduces with *r*_*max*_ in scenarios 1 and 3; this is caused by more frequent acquisition of prey items (as regrowth keeps pace with depletion), which results in a greater frequency of handling rather than foraging.

In scenario 2 however, the frequency of handling is relatively unaffected by increasing *r*_*max*_ (Fig. 6A). The difference between scenarios 2 and 3 has to do with the change in the frequency of kleptoparasitism (Fig. 6B). In scenario 2, kleptoparasitism forms *>* 75% of all activities at low *r*_*max*_, and is much more common than in scenario 3 populations at the same regrowth rate. However, at relatively high *r*_*max*_ (0.03), the fixed kleptoparasitic strategy goes extinct. This is because at high *r*_*max*_, forager-prey encounters are more common than kleptoparasite-handler encounters, in both early (< 10) and later generations (*>* 50). Consequently, kleptoparasites have relatively much lower fitness than foragers, and do not proliferate. Thus at high *r*_*max*_, a scenario 2 population is nearly identical to a scenario 1 population; while some kleptoparasites may be seen in later generations, these occur most likely due to ephemeral mutations in the forager strategy.

**Figure 6:**
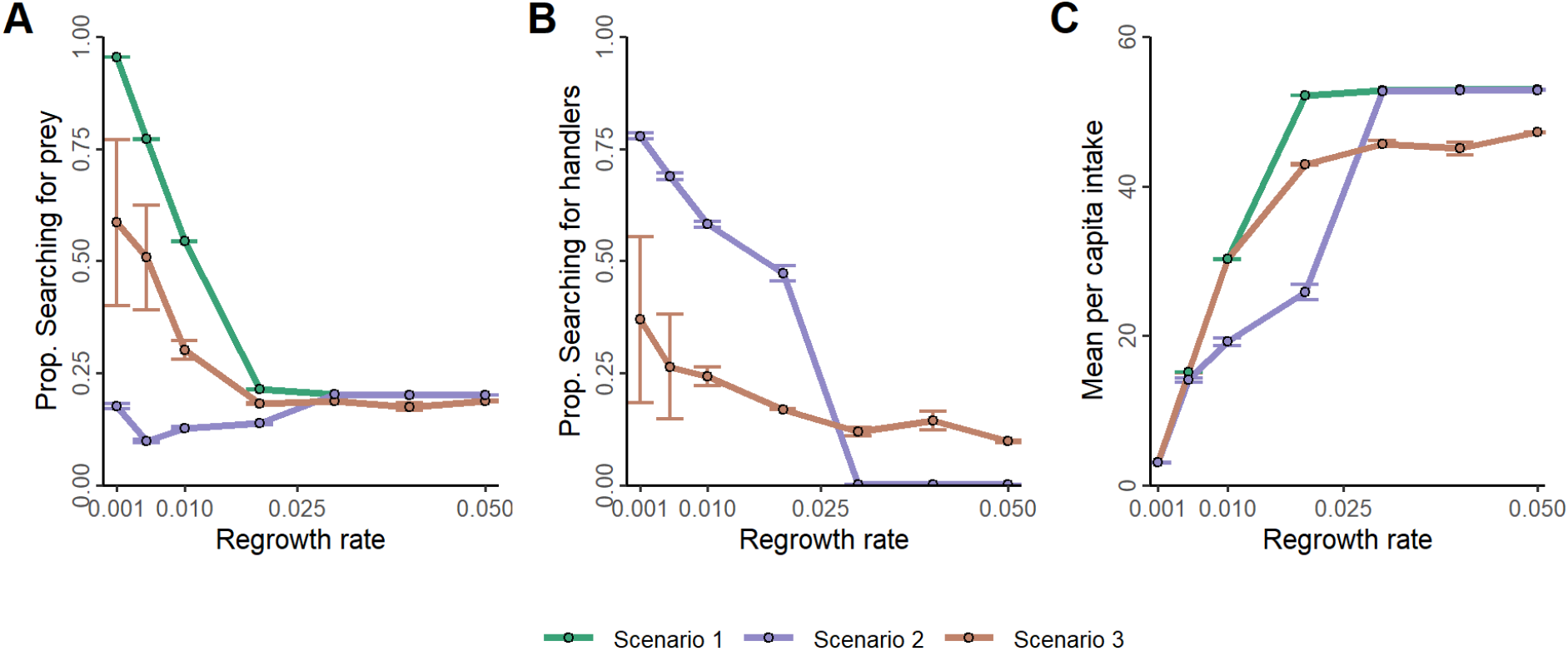
Landscape productivity strongly affects scenario outcomes. **(A)** The proportion of time spent searching for food decreases with increasing *r*_*max*_ in scenarios 1 and 3 but remains relatively stable within scenarios. This is partly due to a higher proportion of time spent handling at higher prey densities. **(B)** The proportion of time spent searching for handlers (in order to steal prey from them) also decreases with increasing *r*_*max*_. In scenario 2, kleptoparasites go extinct for *r*_*max*_ values above 0.025. **(C)** At low productivity, the average intake is similar in all three scenarios. For higher *r*_*max*_ values the average intake rate is lowest in scenario, until *r*_*max*_ is larger than 0.025 and kleptoparasites go extinct (leading to the same kind of population as in scenario 1). At high *r*_*max*_, the average intake rate in populations with conditional kleptoparasites (scenario 3) is substantially lower than in populations without kleptoparasitism.

In scenario 3, kleptoparasitism persists at low frequencies even at the highest regrowth rates (Fig. 6B); thus some foragers lose time in extracting items which are then stolen from them. Consequently, while populations in all three scenarios achieve very similar mean per-capita intakes at low *r*_*max*_, at intermediate regrowth rates (0.01, 0.02), conditionally kleptoparasitic populations achieve a higher mean per-capita intake than populations using fixed strategies. Only at high *r*_*max*_, when fixed strategy populations effectively convert to purely forager populations, do they achieve a higher intake than conditional strategy populations (Fig. 6C).

## Discussion

Our spatially-explicit individual-based model implements the ecology and evolution of movement and foraging decisions, as well as resource dynamics, in biologically plausible ways, and offers a new perspective about the distribution of animals in relation to their resources under different scenarios of competition. First, we show that when moving with a limited perception range and competing only by exploitation, individuals evolve movement strategies for both direct and indirect resource cues (prey items and handlers, respectively). Regardless, on a resource landscape with discrete prey items, large areas may become devoid of any movement cues, leading to a mismatch between individual distribution, prey item distribution, and landscape productivity. Second, we show that when interference competition in the form of kleptoparasitism is allowed as a fixed strategy, it rapidly establishes itself on landscapes where stealing is more time-efficient than searching for prey. This rapid increase in kleptoparasitism as a strategy is accompanied by the evolution of movement strategies that favour moving towards handlers, which are the primary resource of the kleptoparasites. In this sense, obligate kleptoparasites may be thought of as forming a higher trophic level, with any handling consumers as their prey. Third, we show that when foraging strategy is allowed to be conditional on local cues, *(1)* the population’s mean per capita intake is significantly higher than that of a population with fixed strategies, and *(2)* unlike fixed strategy populations, kleptoparasitism as a strategy does not go extinct on high-productivity landscapes. However, across scenarios, individuals are broadly unable to match the productivity of the resource landscape, contrary to the predictions of IFD based models, which predict input matching for some (Parker and Sutherland, 1986; Holmgren, 1995; Hamilton, 2002), or all of the competitive types Korona (1989).

### Comparison with Existing Models

Existing models of competition and movement impose fixed movement rules on individuals to mimic either ideal or non-ideal individuals (Vickery et al., 1991; Cressman and Křivan, 2006; Amano et al., 2006; Beauchamp, 2008; Stillman and Goss-Custard, 2010; White et al., 2018). When individual competitive strategies are included in models, they represent differences in competitive ability (e.g. Parker and Sutherland, 1986; Holmgren, 1995; Hamilton, 2002), or a probabilistic switch between producing and scrounging (Beauchamp, 2008). In contrast, our model allows individuals’ movement (and competition) decisions to be adaptive responses to local environmental cues. Similar to Getz et al. (2015, 2016) and White et al. (2018), our individuals choose from among the available movement options after weighing the local environmental cues, similar to resource selection functions (Manly et al., 2007; White et al., 2018). Local environmental cues in our model are constantly changing, as we model discrete, depletable prey-items, contrasting with many IFD models (Tregenza, 1995; Amano et al., 2006). This allows for a more plausible, fine-scale consideration of exploitation competition, which is often neglected, and allows the cues sensed by individuals to strongly structure the distribution of competitors (see below).

Adaptive responses must have an explicit evolutionary context, and consider multiple generations of the population. We follow Beauchamp (2008) and Getz et al. (2015) in allowing the decision making weights for movement, and variation thereof, to be the outcomes of natural selection. However, instead of using ‘evolutionary algorithms’ (Beauchamp, 2008; Getz et al., 2015, 2016) to ‘optimise’ individual movement rules, we consider a more plausible evolutionary process: Instead of allowing the fittest 50% of the population to replicate, the number of offspring are proportional to individual fitness. The weight loci are subject to mutations independently, rather than subjecting all loci of an individual to simultaneous mutation. Finally, we avoided the unrealistic assumption of ‘simulated annealing’, which adapts the mutation rate or the mutational step sizes to the rate of evolutionary change. Instead we drew mutation sizes from a Cauchy distribution, so that most mutations are very small, but large-effect mutations do occur throughout the simulation. Similarly, rather than determining competition strategy probabilistically or ideally (Vickery et al., 1991; Beauchamp, 2008; Tania et al., 2012), our individuals’ competition decisions are also shaped by selection (in scenarios 2 and 3).

### Movement Rules on Depleted Landscapes

In scenario 1, depletion of discrete prey can leave many areas empty of prey-items: in such areas, movement informed by a resource gradient is impossible, and individuals may move randomly (Perkins, 1992). This lack of direct resource cues for locally optimal movement might be among the mechanisms by which unsuitable ‘matrix’ habitats modify animal movement on heterogeneous landscapes (Kuefler et al., 2010). When individuals do not sense resource gradients, the presence of more successful conspecifics may indicate a suitable foraging spot (local enhancement; Giraldeau and Beauchamp, 1999; Beauchamp, 2008; Cortés-Avizanda et al., 2014). The presence of unsuccessful individuals, meanwhile, may signal potential costs from exploitation or interference competition. This selects for movement strategies incorporating the presence and condition of competitors into individual movement decisions (‘social information’: Dall et al., 2005). Consequently, consumer aggregation — often explained by invoking external costs such as predation (Krause and Ruxton, 2002; Folmer et al., 2012) — could also be the outcome of movement rules that have evolved to trade competition costs for valuable social information on the underlying drivers of the spatial structure (here, *r*) of uninformative landscapes (Folmer et al., 2010; Cortés-Avizanda et al., 2014).

### Individual Variation in Movement Rules

We find substantial individual variation in the strength of movement weights within populations, as expected from heterogeneous landscapes (see Supplementary Material Fig. 2.1 – 2.3; see Wolf and Weissing 2010 for background). The persistence of multiple ‘movement morphs’ across generations indicates that they are alternative movement strategies of equal fitness (see Getz et al., 2015). Indeed, polymorphism in movement rules may help reduce competition as individuals make subtly different movement and competition decisions when presented with the same cues (Laskowski and Bell, 2013, see also Wolf and Weissing 2012). Scenario 2 also shows significant within-strategy individual variation in movement weights, which might ameliorate within-strategy exploitation competition, or help foragers avoid kleptoparasites (Wolf and Weissing, 2012; Laskowski and Bell, 2013). Interestingly, scenario 3 has the least individual variation in movement rules, potentially because plasticity in competition strategy dampens such diversification (Pfennig et al., 2010), but also possibly because the ability to switch between prey types reduces the intensity of competition. Here, non-handler avoidance shows the most morphs, but it is unclear whether this variation is linked to the frequency with which individuals use either foraging strategy — potentially leading to subtle, emergent behavioural differences that are conditioned on the local environment (Wolf and Weissing, 2010, 2012).

### Competition Strategies and the IFD

IFD models predict that individual movement should result in consumer distributions tracking the profitability of resource patches (Fretwell and Lucas, 1970; Parker, 1978), with dominant competitive types (including kleptoparasites) monopolising the best patches (Parker and Sutherland, 1986; Holmgren, 1995; Hamilton, 2002, but see Korona 1989). In scenarios 2 and 3, kleptoparasitic individuals unsurprisingly and rapidly evolve to track handlers (a direct resource), while avoiding non-handlers (potential competitors). However, these evolved rules do not lead kleptoparasites to occupy the best cells as predicted (Parker and Sutherland, 1986; Holmgren, 1995; Hamilton, 2002). Across our scenarios (including scenario 1), individual density is only weakly correlated with cell productivity. In scenario 2, this departure from predictions is driven by the contrasting movement rules of foragers, which evolve to *avoid* handlers as well as non-handlers, both of which might be kleptoparasites (cryptic interference; seen in interference-sensitive waders Folmer et al. 2010; Bijleveld et al. 2012; see Supplementary Material). Thus, foragers likely avoid resource peaks, which are more likely to have handlers (due to the higher probability of forager-prey encounters Parker and Sutherland, 1986; Holmgren, 1995; Hamilton, 2002). Fixed kleptoparasites cannot extract prey themselves, and must move off resource peaks to track and rob handlers (similar to Parker and Sutherland, 1986), breaking the link between individual density and productivity. This shows the pitfalls of simplistically linking current ecological conditions with population distributions without considering competitive strategies or evolutionary history.

### Constraints on Competition Strategies

Foraging strategies involving specialisation on a resource type are expected to be constrained by the availability of that resource; thus kleptoparasitism, seen as a prey-choice problem, should be constrained by the density of targets (Ens et al., 1990). In scenarios 2 and 3, more kleptoparasitism should be expected with increasing *r*_*max*_, as prey and consequently, handlers, are expected to be more abundant. Instead, kleptoparasitism declines with increasing *r*_*max*_, in line with Emlen (1966), who predicted that the commoner food type (prey) rather than the more efficiently exploited one (handlers) should be preferred. This effect is especially stark in scenario 2, where kleptoparasites go extinct when prey are very common at high *r*_*max*_. At stable population densities, the persistence of fixed kleptoparasitism depends on their intake *relative to foragers*. Since intake is an outcome of movement rules, and population movement rules are not well adapted to the environment in early generations, foragers obtain, as a clade, more intake than kleptoparasites. Modelling discrete prey-items and individuals in a spatial context, then, leads to the finding that obligate kleptoparasitism is only a viable strategy when forager-prey encounters are less common than kleptoparasite-handler encounters. This might explain why — and is supported by the observation that — kleptoparasitism is common among seabirds, whose communal roosts are much easier targets than unpredictable shoals of fish out at sea (Brockmann and Barnard, 1979); in contrast, grazing geese have similar flock sizes but their resource is also very easily located, hence kleptoparasitism is rare even though interference is common (Amano et al., 2006). Finally, comparing across regrowth rates shows why possibly cryptic behavioral complexity should be considered in predictions of the long-term effect of environmental change on populations. While both scenario 1 and 2 populations appear identical at high *r*_*max*_, even a small decrease in environmental productivity could lead to an abrupt drop in per-capita intake — and potentially, strongly reduced growth or survival — for fixed strategy populations due to unexpected, emergent kleptoparasitism.

### Comparison with Conceptual Models

Classical models of animal movement and foraging largely consider homogeneous populations and environmental conditions, and movements that are made either optimally or at random. While these models provide powerful insights, individual-based models such as ours have the advantage that they can accommodate individual variation, local environmental conditions, and the mechanisms of movement and decision-making. Individual-based modeling has the obvious drawback that numerous specific assumptions have to be made, which might not all be founded on empirical evidence, and might seem to limit the generality of the conclusions. Nevertheless, as long as these models are not mistaken for attempts at faithful representations of real systems, their exploration provides valuable perspectives on the conceptual models that have dominated theory in the past. After all, traditional models also include numerous assumptions (the spatiotemporal structure, the timing of events, the distribution and inheritance of traits) that are usually not stated and therefore less visible. For the future, we envisage pluralistic approaches, where both types of model are applied to the same research question. Only comparing the outcomes of diverse models will reveal which conclusions and insights are robust, and which reflect peculiarities of the model structure Only such model comparison can tell us whether and when simple models produce general insights, where simple models fail, and when mechanisms can explain initially counterintuitive observations, such as the attraction to competitors that we observed in our study.

### Individual Based Models in Movement Ecology

Animal movement ecology takes an explicitly individual-based approach, centred around individual decisions (Nathan et al., 2008). This makes individual-based models a reasonable choice when seeking general insights into the evolutionary ecology of animal movement strategies (see e.g. Getz et al., 2015), whose ultimate causes are otherwise difficult to study empirically. They can incorporate local circumstances and state variables in considerable detail, and thereby promote careful consideration of what we know about animal response mechanisms. Individual-based models of movement decisions can also be related to existing empirical work in animal tracking. For example, our model’s decision making weights are likely familiar to movement ecologists in the form of the individual-specific coefficients of resource-selection or step-selection functions, and have been interpreted as such (White et al., 2018). By allowing selection coefficients from animal-tracking studies to undergo natural selection on simulated landscapes, similar models could help explore long-term changes in movement strategies. This approach would require very accurate estimation of the fitness outcomes of movement — no easy task. Consequently, individual-based models are not (yet) intended to be ‘fit’ to empirical movement data. Rather, they are useful to elucidate how simple mechanisms can lead to unexpectedly complex outcomes, and to help define a broad envelope of potential outcomes, given known mechanisms and explicit assumptions. These outcomes can provide valuable perspective on population-level models (such as the IFD), or be used to explore how movement strategies evolve in dynamic environments.

## Data and Code Availability

Simulation model code is on Github: github.com/pratikunterwegs/Kleptomove and Zenodo: zenodo.org/record/4905476. Simulation data are available from DataverseNL as a draft: https://dataverse.nl/privateurl.xhtml?token=1467641e-2c30-486b-a059-1e37be815b7c. Data will be at this persistent link after publication: doi.org/10.34894/JFSC41. Data analysis code is on Github: github.com/pratikunterwegs/kleptomove-ms and on Zenodo: doi.org/10.5281/zenodo.4904497.

## Acknowledgments

The authors thank Hanno Hildenbrandt for contributing extensively to the coding of the simulation model *Kleptomove*; Matteo Pederboni for contributing to the model’s development; and members of the Modelling Adaptive Response Mechanisms Group, and of the Theoretical Biology department at the University of Groningen for helpful discussions on the manuscript. F.J.W. and C.F.G.N. acknowledge funding from the European Research Council (ERC Advanced Grant No. 789240). P.R.G was supported by an Adaptive Life Programme grant made possible by the Groningen Institute for Evolutionary Life Sciences (GELIFES).

## Notes

https://github.com/pratikunterwegs/kleptomove-ms

https://github.com/pratikunterwegs/Kleptomove

https://zenodo.org/record/4905476

https://zenodo.org/record/5112915

https://dataverse.nl/privateurl.xhtml?token=1467641e-2c30-486b-a059-1e37be815b7c

### Competing Interest Statement

The authors have declared no competing interest.

### Summary of Updates

Revised for length.

