## Supplementary Material for "The joint evolution of animal movement and competition strategies"

2021-07-19

In this Supplementary Material, we show two separate kinds of figures.

In Section 1, **Landscape Depletion across $r_{max}$**, we show the development of the prey-item landscape across different regrowth rates that we explored for our model. A summary of the eco-evolutionary dynamics for each $r_{max}$, in each scenario, can be found in the main text (see Main Text Figure 6).

In Section 2, **Evolution of Decision Making Weights**, we show the frequency of the movement decision making weights across generations, for an $r_{max}$ of 0.01 (the default value, with results presented in the main text). In addition, for scenario 3, we show the evolution of the strategy weight with respect to handlers, i.e., the foraging strategy response of individuals to the presence of handlers.

### 1 Landscape Depletion across $r_{max}$

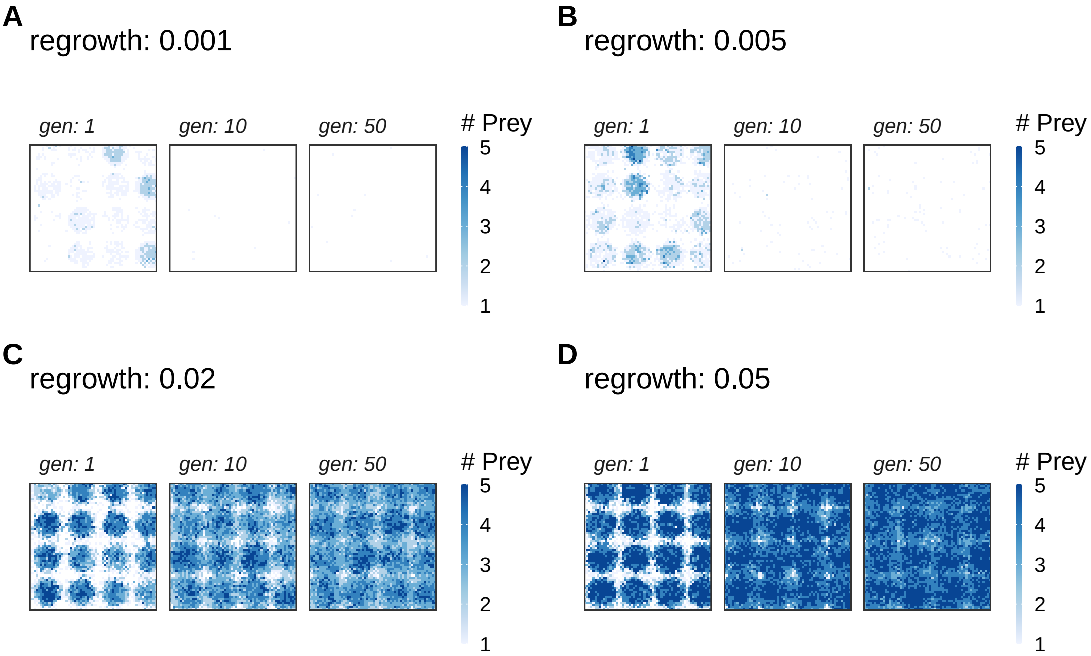


Figure 1.1: In scenario 1, foragers completely deplete the resource landscape within 10 generations at low $r_{max}$ (A, B). However, at $r_{max} > 0.01$ (C, D), prey item regeneration exceeds depletion and the resource landscape is rapidly saturated until most cells carry 5 items, the maximum allowed in our model.

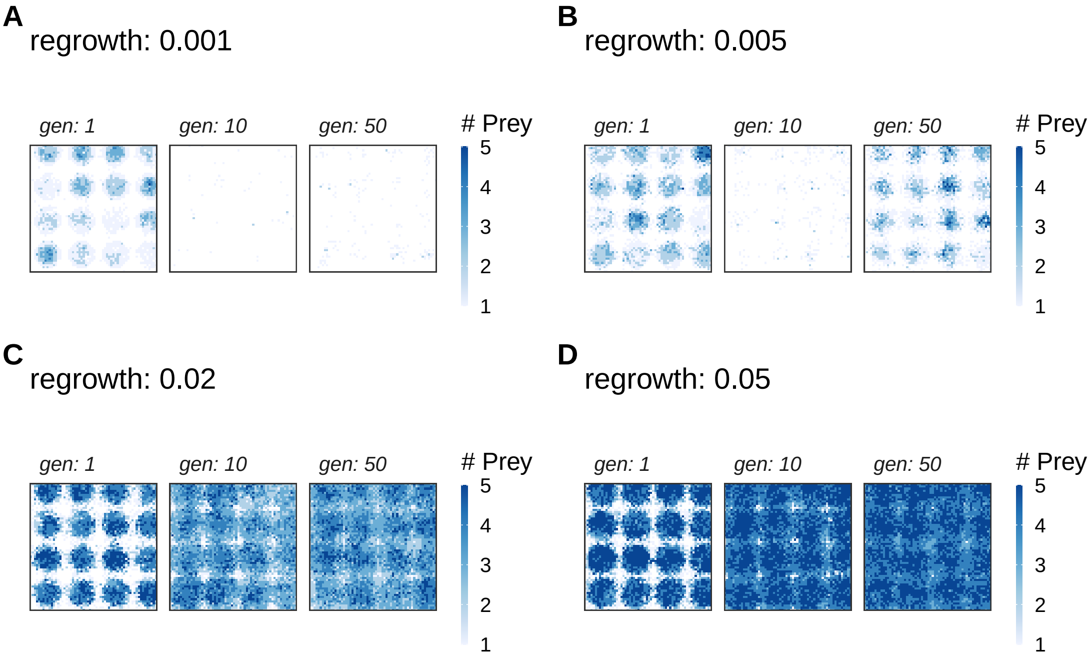


Figure 1.2: In scenario 2, foragers can only deplete the resource landscape at very low $r_{max}$ (A): 1 prey item generated per 1,000 timesteps, or 2.5 generations. At all $r_{max} \geq 0.05$ (B, C, D), prey item regeneration matches or exceeds depletion and the resource landscape either shows strong spatial structure, or is entirely saturated with prey items.

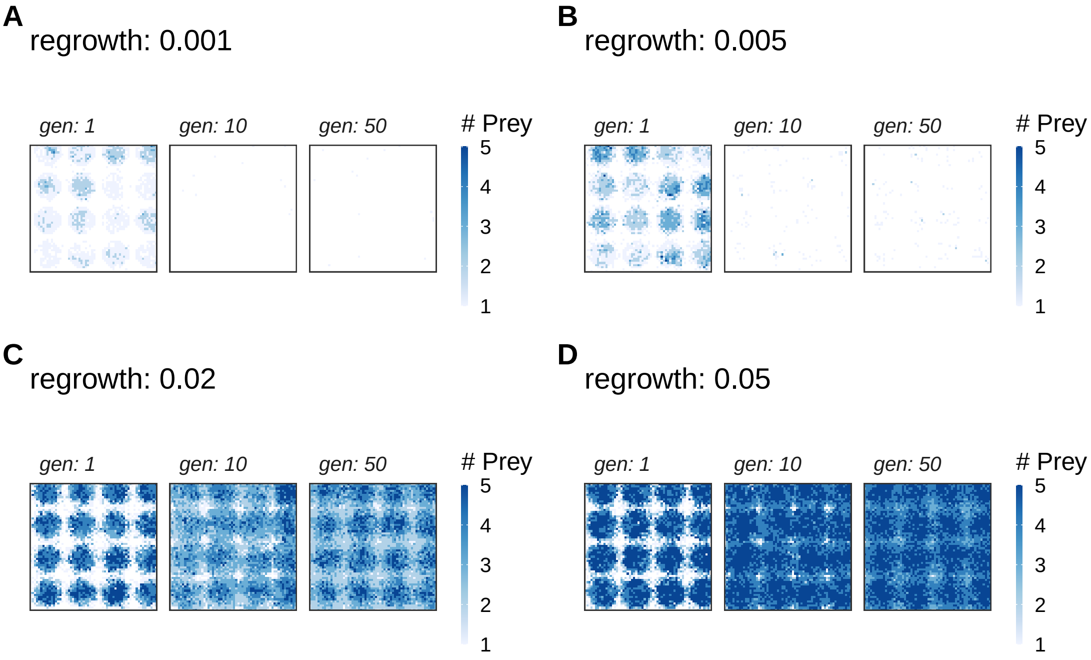


Figure 1.3: Scenario 3 is similar to scenario 1 at low $r_{max}$ (A, B), where foragers completely deplete the resource landscape). Similarly, at $r_{max} > 0.01$ (C, D), prey item regeneration exceeds depletion and the resource landscape is rapidly saturated to a carrying capacity of 5 prey items per cell.

### 2 Evolution of Decision Making Weights

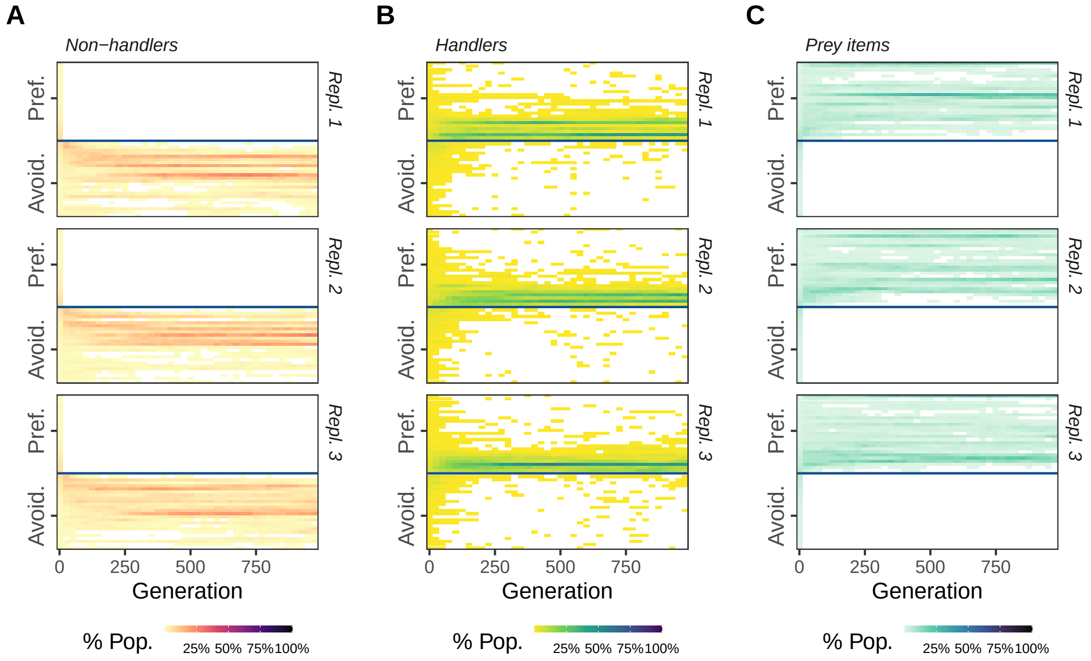


Figure 2.1: In scenario 1, populations evolve (A) to move away from non-handlers, (B) move towards handlers, and (C) to move towards cells with more prey items. While the sign of the response (avoidance or preference) is consistent across replicates, the replicates differ in the number and frequency of evolved responses (i.e., decision-making weight values). All panels show an $r_{max} = 0.01$.

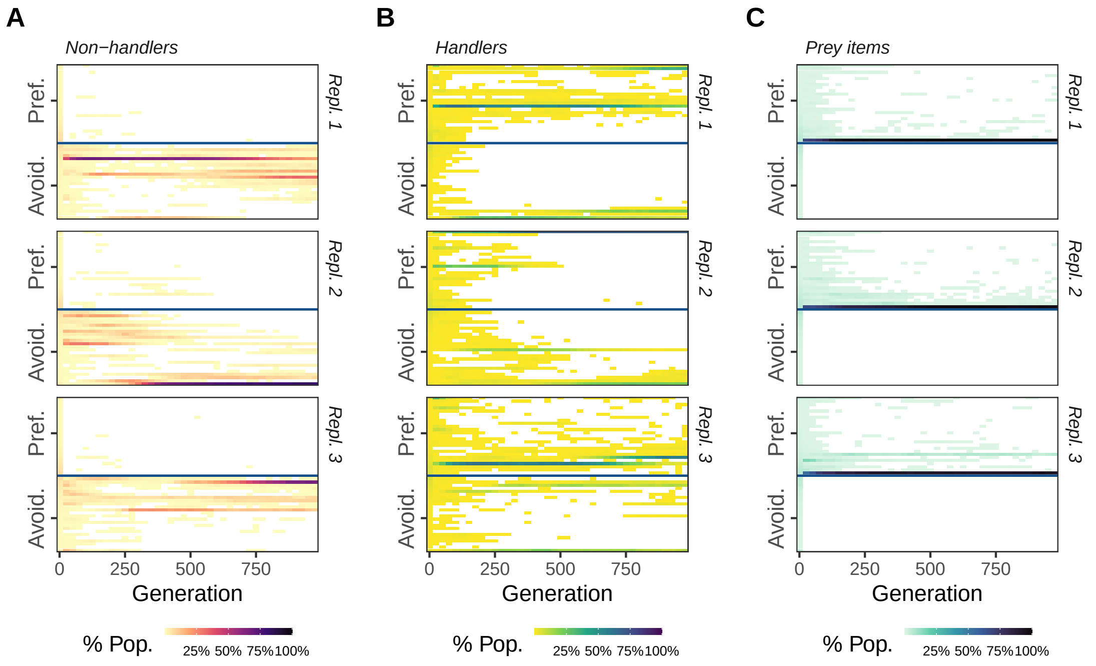


Figure 2.2: In scenario 2, populations evolve (A) to move away from non-handlers, but (B) a mixed response towards handlers due to the correlation between handler-preference and the kleptoparasitic strategy (see Fig. 3, main text). Here too, responses are polymorphic, with little consistency across replicates, despite the overall sign of the response being consistent. (C) In contrast with scenario 1, most foragers show only a weak preference for moving towards cells with more prey items. All panels show an $r_{max} = 0.01$.

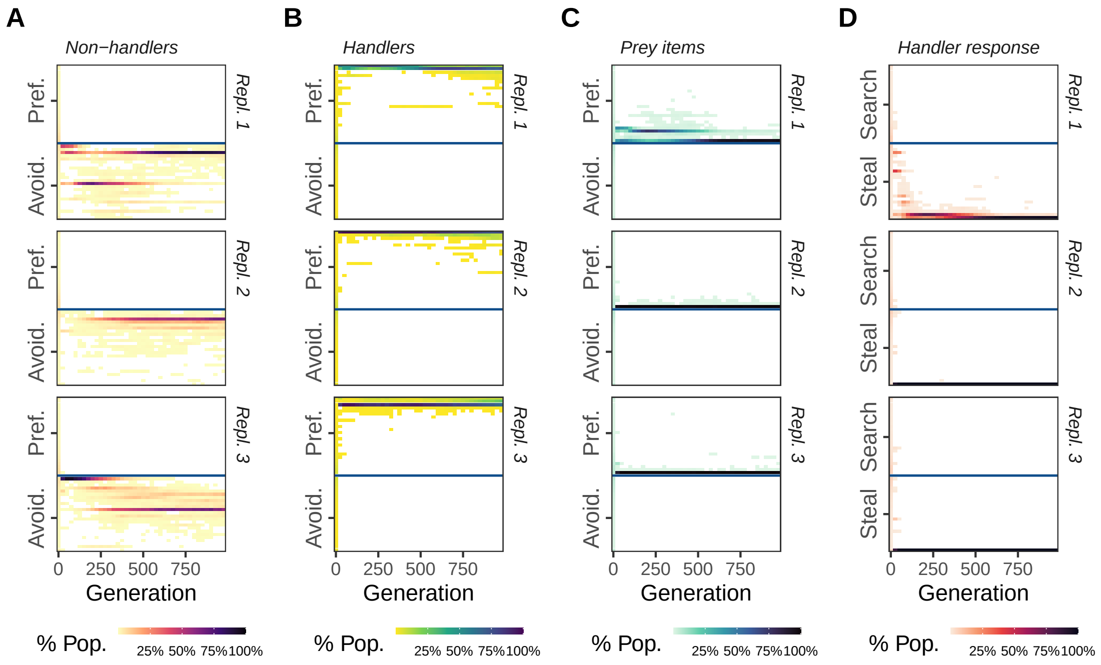


Figure 2.3: In scenario 3, populations evolve (A) to move away from non-handlers, and (B) towards handlers. (C) Most foragers also show a weak preference for moving towards cells with more prey items. (D) All individuals show a kleptoparasitic response to handlers. All panels show an $r_{max} = 0.01$.
